## Supplementary file 1 for "Operation regimes of spinal circuits controlling locomotion and role of supraspinal drives and sensory feedback"

**Supplementary file 1: Sensitivity analysis of a single rhythm generator model to changes of its key parameters**

We studied the sensitivity of the model of a single rhythm generator (RG) to variations of key parameters by analyzing the effects of these variations on (1) changes in the durations of flexor and extensor activity as a function of the external drive to the F half-center (Drive-F) and (2) the values of Drive-F at which the RG model switches from the state-machine regime to the flexor-driven rhythm generation regime (transition 1) and from the frexor-driven rhythm generation regime to the classical half-center regime (transition 2). Transition 1 was defined by the value of Drive-F = $DF_{T1}$, at which the flexor half-center started generating intrinsic oscillations. Transition 2 was defined by the value of Drive-F = $DF_{T2}$, when the duty factor (extensor phase/period of oscillations) became equal to 0.5, after which the length of flexor bursts started to increase with the increasing Drive-F.

Our simulations have demonstrated that both $DF_{T1}$ and $DF_{T2}$ depend on the expression (maximal conductance) of the persistent sodium current ($\bar{g}_{\mathrm{NaP}}$), the excitability of F and E half-centers, and the strength of mutual inhibition between them. The excitability of half-centers is defined by the parameters of leakage current (the leakage conductance $g_{\mathrm{L}}$ and the leakage reversal potential $E_{\mathrm{L}}$) and excitatory drives to them (Drive-F and Drive-E). The mutual inhibition between F and E centers depends on the weights of inhibitory connections between the centers. Therefore these parameters were selected to study sensitivity of the RG model. We focused on the effects of parameter variations on the durations of F and E phases and the values of $DF_{T1}$ and $DF_{T2}$, critical time moments defining the transitions between the RG regimes. In this analysis, we used OFAT (One-factor-at-a-time) method (Daniel H One-at-a-time plans. J. Am. Stat. Association **68:**353-360 (1973).

Figure 1S shows the results of changes in the duration of F and E phases, and $DF_{T1}$ and $DF_{T2}$ values based on variation of the following key parameters: $\bar{g}_{\mathrm{NaP}}$ (panels A1-A3), $g_{\mathrm{L}}$ (panels B1-B3), $E_{\mathrm{L}}$ (panels C1-C3), the weights of inhibitory connections from InF to E ($w_4$, panels D1-D3) and from InE to F neurons ($w_2$, panels E1-E3), and constant drive to the E half-center, Drive-E (panels F1-F3).

In most cases, these manipulations produced moderate to small opposite changes in the durations of F and E phases. The strongest effects were produced by the variations of $\bar{g}_{\mathrm{NaP}}$ (panels A1 and A2,) and $g_{\mathrm{L}}$ (panels B1 and B2, especially B1). Elevation of $\bar{g}_{\mathrm{NaP}}$ (panel A3) or reduction of $g_{\mathrm{L}}$ (panel B3) shifted $DF_{T1}$ and $DF_{T2}$ points to lower values of Drive-F. Similar, but weaker effects were produced by increasing the negative value of $E_{\mathrm{L}}$ (panels C1-C3) or by a reduction of $w_2$ (panels E1-E3) and Drive-E (panels F1-F3). Reduction of $w_4$ (panels E1-E3) affected only the duration of the F phase.

We concluded that the duration of F and E phases and the values of $DF_{T1}$ and $DF_{T2}$, defining the transition between RG regimes, were most sensitive to variations of the intrinsic parameters of the RG half-centers: $\bar{g}_{\mathrm{NaP}}$ and $g_{\mathrm{L}}$.

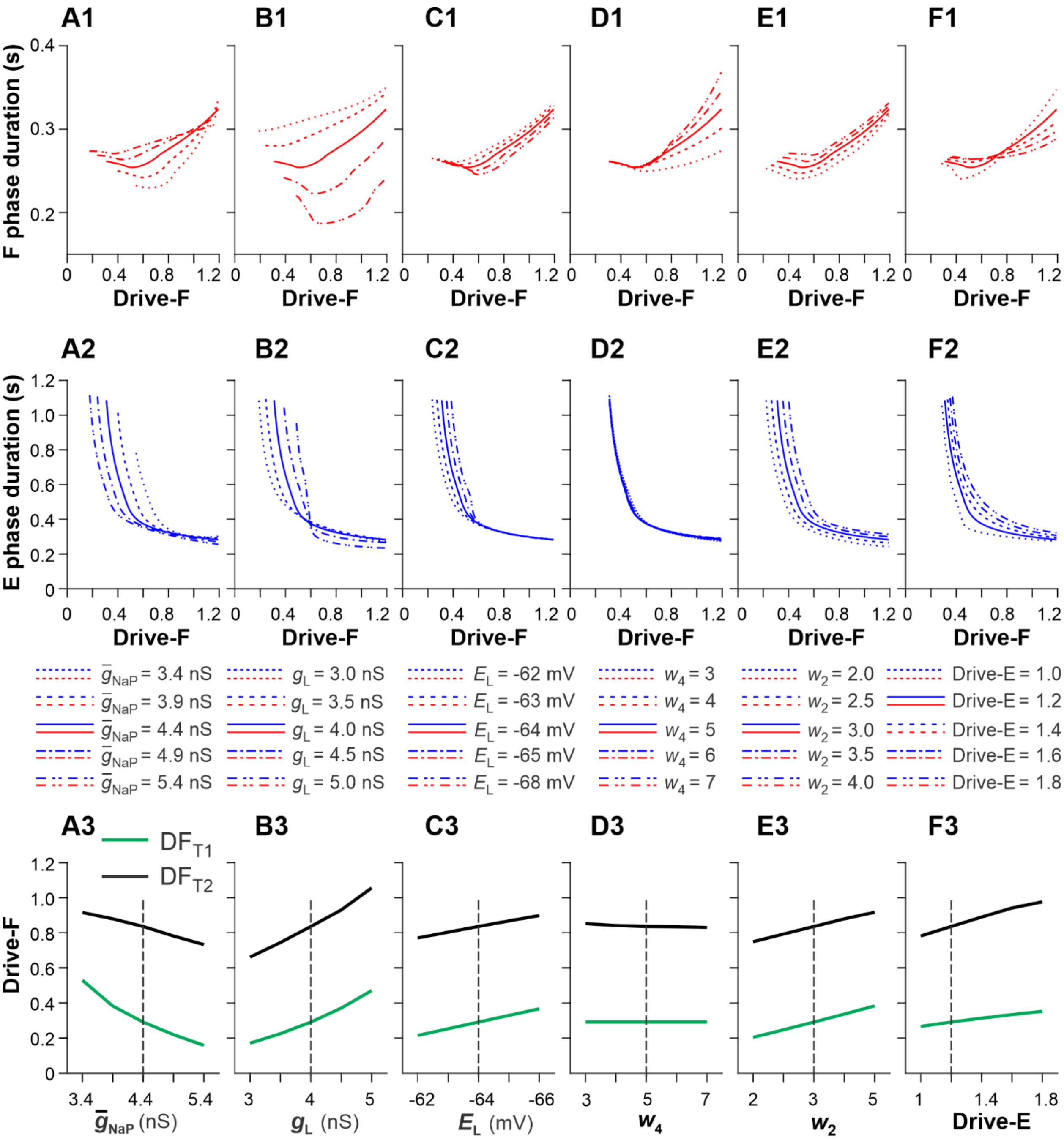


**Figure 1S. Model sensitivity (F and E phase durations and** $DF_{T1}$ and $DF_{T2}$ **values) to variations of key model parameters:** $\bar{g}_{NaP}$ (**A1-A3**), $g_L$ (**B1-B3**), $E_L$ (**C1-C3**), the weights of inhibitory connections from InF to E ($w_4$, **D1-D3**) and from InE to F ($w_2$, **E1-E3**), and Drive-E (**F1-F3**). Bold lines in the two upper rows and vertical dashed lines in the bottom row correspond to the basic values of model parameters.
